# Dependency of Lower Limb Joint Reaction Forces on Femoral Anteversion

**DOI:** 10.1101/2021.02.22.432159

**Authors:** Luca Modenese, Martina Barzan, Christopher P Carty

## Abstract

**Background:** Musculoskeletal (MSK) models based on literature data are meant to represent a generic anatomy and are a popular tool employed by biomechanists to estimate the internal loads occurring in the lower limb joints, such as joint reaction forces (JRFs). However, since these models are normally just linearly scaled to an individual’s anthropometry, it is unclear how their estimations would be affected by the personalization of key features of the MSK anatomy, one of which is the femoral anteversion angle.

**Research Question:** How are the lower limb JRF magnitudes computed through a generic MSK model affected by changes in the femoral anteversion?

**Methods:** We developed a bone-deformation tool in MATLAB (https://simtk.org/projects/bone_deformity) and used it to create a set of seven OpenSim models spanning from 2° femoral retroversion to 40° anteversion. We used these models to simulate the gait of an elderly individual with an instrumented prosthesis implanted at their knee joint (5^th^ Grand Challenge dataset) and quantified both the changes in JRFs magnitude due to varying the skeletal anatomy and their accuracy against the correspondent in vivo measurements at the knee joint.

**Results:** Hip and knee JRF magnitudes were affected by the femoral anteversion with variations from the unmodified generic model up to 11.7±5.5% at the hip and 42.6±31.0% at the knee joint. The ankle joint was unaffected by the femoral geometry. The MSK models providing the most accurate knee JRFs (root mean squared error: 0.370±0.069 body weight, coefficient of determination: 0.764±0.104, largest peak error: 0.36±0.16 body weight) were those with the femoral anteversion angle closer to that measured on the segmented bone of the individual.

**Significance:** Femoral anteversion substantially affects hip and knee JRFs estimated with generic MSK models, suggesting that personalizing key MSK anatomical features might be necessary for accurate estimation of JRFs with these models.

## Introduction

Computational models of the musculoskeletal (MSK) system derived from cadaveric studies or literature data, also known as *generic models*, are commonly employed to estimate internal joint forces during healthy and pathological gait. The underlying assumption is that their “average” MSK anatomy can be scaled to produce a satisfactory representation of an individual’s lower limb. Indeed, generic models have demonstrated remarkable accuracy in estimating lower limb joint loadings when compared to in vivo measurements from instrumented prostheses [1, 2], both at the hip [3] and at the tibiofemoral joint [4]. Moreover, compared to image-based subject-specific models, which are time consuming and technically challenging to generate [5], generic models are straightforward to scale and use in biomechanical workflows, and represent a reliable alternative to traditional direct kinematic models [6]. Nonetheless, the uptake of MSK modelling for the computation of kinematics, kinetics and joint forces in hospital-based gait laboratories is sparse, due in part to the absence of editing tools to modify key anatomical features, such as the lower limb torsional profile, in existing MSK modelling frameworks. A systematic quantification of the impact of this lack of personalization on the accuracy of joint reaction forces (JRFs) is unavailable in previous MSK modelling literature [7]. For example, femoral anteversion, the angle between the femoral neck axis and the posterior condylar axis, has been previously shown to affect muscle moment arms in children affected by cerebral palsy [8] and to influence the hip JRFs in hip replacement patients [9] and typically developing children [10], however, its effect on the knee and ankle joints remains unknown.

In this work we will contribute to the state of the art by presenting a tool for applying user-defined torsional lower-limb profiles to generic musculoskeletal models in OpenSim and investigating the dependency of the JRF magnitudes to varying femoral anteversion angles in a scaled generic MSK model of the lower limb.

## Materials and methods

### Bone-deformation tool

A bone-deformation toolbox for MSK models was implemented in MATLAB using the Application Programming Interface of OpenSim 3.3 [11]. This tool can apply any linear user-defined torsion profile to the long axis of a specified bone (Figure 1A), generating a deformed skeletal anatomy and adjusting the muscle attachments accordingly (Figure 1B). At the user discretion, the joint parameters can be modified together with the musculoskeletal anatomy, so enabling both pure bone morphological alterations and the modelling of deformities affecting the lower limb kinematics. Here we will focus on the former functionality, but examples of the latter are presented in the supplementary materials.

**Figure 1.**
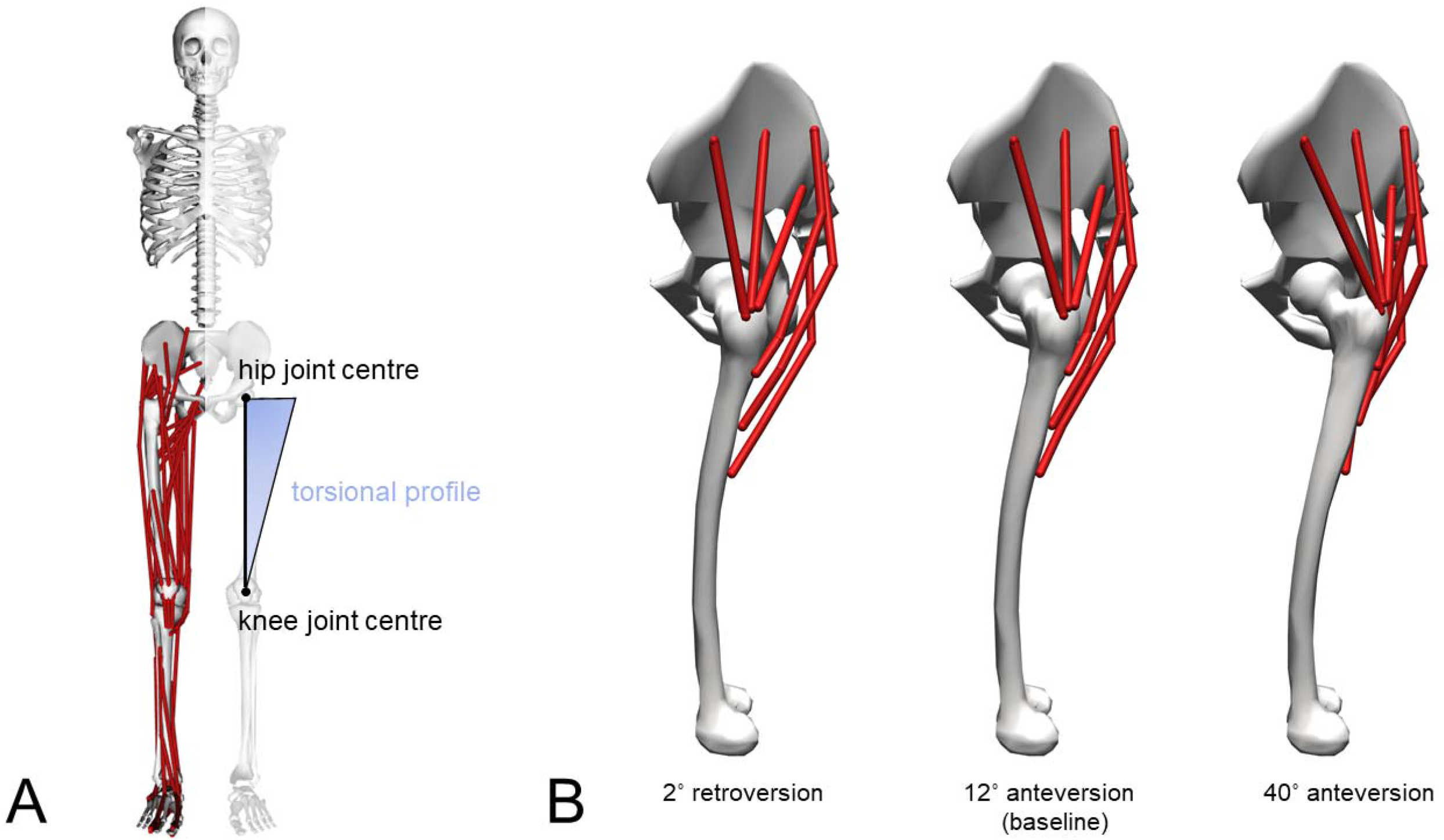
(A) The musculoskeletal anatomy used as baseline in this study and a schematic representation of the torsion profile employed to generate the modified models, represented on the left leg. The pelvis and modified femur of the models with minimum (2° retroversion) and maximum (40° anteversion) anteversion angles, together with the baseline model, are shown together with the gluteus medius and gluteus maximus geometries (B).

The bone-deformation tool can be downloaded from https://simtk.org/projects/bone_deformity and is openly developed at https://github.com/modenaxe/msk_bone_deformation.

### Baseline model and deformed models

The generic full-body model of Rajagopal et al. [12] was first modified by removing the upper limbs and then linearly scaled to the anthropometrics of an elderly individual with an instrumented total knee prosthesis implanted on his left leg (age: 86, mass: 75 kg, data from the 5^th^ Grand Challenge dataset shared at https://simtk.org/projects/kneeloads [2]). The model was further adjusted by decreasing the maximum isometric forces of the muscles crossing the knee joint by 40% to model the decrease in joint strength following knee replacement, similarly to [13], so obtaining the *baseline model*. Subsequently, the left femur geometry (estimated anteversion angle: 12°, see supplementary materials) was altered using the bone-deformation MATLAB tool. Six *modified models* were generated with angles from 2° retroversion to 40° anteversion in 7° steps, reflecting the range reported by Strecker et al. [14]. Apart from the left femur alterations, the baseline and modified models were identical to decouple the effects of the bone alteration from the other model features.

### Simulations

Using a standard workflow (inverse kinematics, static optimization with quadratic objective function, joint reaction analysis) in OpenSim 3.3, five level walking trials performed at self-selected speed were simulated with all the models, calculating the JRF peak magnitudes for the left leg joints. These values were then compared among baseline and modified models to quantify their percentage peak variations and, at the knee joint, their accuracy against the correspondent in vivo measurements using peak errors (normalized to body weight, BW), root mean squared errors (RMSE) and coefficients of determination R^2^.

## Results

The computed JRF magnitudes showed a marked dependency on femoral anteversion (Figure 2, Table 1) at the hip and knee joints, while the ankle joint loading was practically unaffected (<1% differences). The JRF variations increased with the anteversion angle, up to 11.7±5.5% of the baseline model values for the hip joint and 42.6±31.0% for the knee joint. These loadings decreased for lower-than-baseline-anteversion up to −7.45±1.46% and −2.0±2.33% for the hip and knee joint respectively.

**Figure 2.**
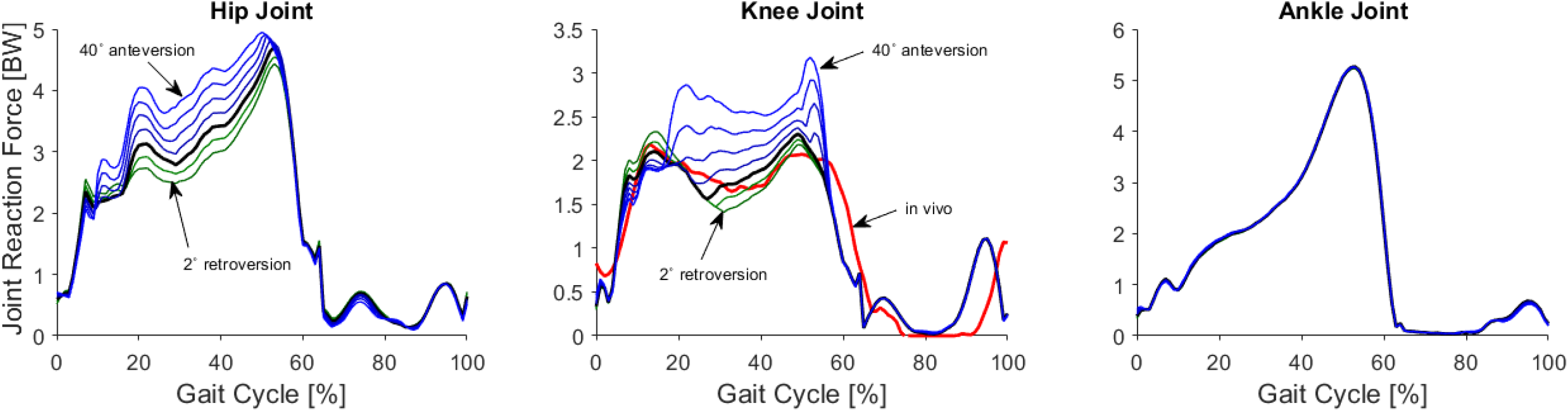
Joint reaction forces computed for a representative gait trial (“PS_ngait_og_ss1”) at the hip, knee and ankle joints for the baseline musculoskeletal model (black solid line, femoral anteversion angle: 12°) and the modified models. Results from models with increased femoral anteversion (from 19° to 40° anteversion angles, blue lines) and decreased femoral anteversion (from 5° anteversion to 2° retroversion angle, green lines) are plotted for lower limb joints (curves overlap for the ankle joint). In vivo loading from in vivo synchronous measurements is also plotted for the knee joint (red solid line).

**Table 1.** Variations of predicted joint reaction forces in the models with modified femurs and comparisons of knee joint reaction forces against the in vivo knee contact forces recorded by Fregly et al. (2012) for the same walking trials.

| Anteversion Angle | JRF peak variations from baseline model [%] |  |  |  |  |  | Comparison with in vivo knee contact forces |  |  |  |  |  |  |  |
| --- | --- | --- | --- | --- | --- | --- | --- | --- | --- | --- | --- | --- | --- | --- |
|  | hip |  | knee |  | ankle |  | Error at peak 1 [BW] |  | Error at peak 2 [BW] |  | RMSE [BW] |  | R<sup>2</sup> |  |
|  | Mean | std | Mean | std | Mean | std | Mean | std | Mean | std | Mean | std | Mean | std |
| -2* | -7.5 | 1.5 | -0.1 | 4.6 | -0.4 | 0.2 | 0.26 | 0.50 | 0.29 | 0.16 | 0.377 | 0.071 | 0.755 | 0.110 |
| 5 | -3.9 | 0.7 | -2.0 | 2.3 | -0.2 | 0.2 | 0.17 | 0.53 | 0.36 | 0.16 | 0.370 | 0.069 | 0.764 | 0.104 |
| 12 | 0.0 | 0.0 | 0.0 | 0.0 | 0.0 | 0.0 | 0.09 | 0.55 | 0.44 | 0.17 | 0.372 | 0.068 | 0.761 | 0.104 |
| 19 | 4.0 | 0.9 | 5.4 | 6.0 | -0.1 | 0.3 | 0.02 | 0.56 | 0.60 | 0.23 | 0.394 | 0.066 | 0.734 | 0.103 |
| 26 | 7.3 | 2.2 | 17.5 | 15.0 | -0.3 | 0.6 | 0.06 | 0.52 | 0.92 | 0.40 | 0.456 | 0.061 | 0.649 | 0.101 |
| 33 | 9.8 | 4.1 | 30.1 | 24.0 | -0.4 | 0.6 | 0.34 | 0.36 | 1.28 | 0.58 | 0.560 | 0.063 | 0.476 | 0.113 |
| 40 | 11.7 | 5.5 | 42.6 | 31.0 | -0.4 | 0.4 | 0.67 | 0.19 | 1.66 | 0.64 | 0.697 | 0.063 | 0.188 | 0.148 |

\* negative anteversion in this table indicates retroversion.

Based on RMSE and R^2^ values (Table 1), the knee JRFs closer to in vivo measurements were those of the models with 5° and 12° anteversion, which also exhibited the lowest mean errors at the second force peak, 0.36±0.16 BW and 0.44±0.17 BW respectively.

## Discussion

In this work we presented a tool for altering the geometry of long bones in generic MSK models and used it to investigate the effect of femoral anteversion on the JRF magnitudes in the lower limb. Using a full-body MSK model as baseline, we created models representing the distribution of femoral torsion reported in [14] and used the 5^th^ Grand Challenge dataset for evaluating their accuracy. In all models, the maximum isometric force of the knee-spanning muscles was further decreased (40%) compared to Marra et al. [13] (35%) to account for baseline muscle parameters representative of a young individual.

The hip JRFs trend monotonically increasing with anteversion angle is consistent with those reported by Kainz et al. [10] using a commercial bone-deformation tool and by Heller et al. [9] using a different MSK modelling approach. Our hip JRF magnitudes are comparable to [5, 10] and, as in those studies, they are larger than in vivo measurements [1]. In previous literature this overestimation has been attributed to the simplified anatomical representation of the hip muscle anatomy [15]. The adopted model, however, explicitly includes a patellar tendon and patellofemoral joint, and therefore we preferred it to other models as it provides a more realistic baseline for tibiofemoral JRF estimation. At the knee joint, the most accurate JRFs were estimated by the baseline model and the modified model with 5° of femoral anteversion, which presented the closest femoral geometry to the actual participant’s femoral anteversion, estimated to be around 10° using the segmented geometry provided with the Grand Challenge dataset (see supplementary materials).

Our findings suggest that personalized femoral anteversion could improve the JRF estimation at the knee joint. Additionally, given the observed influence of a single morphological feature on the JRF outputs, caution should be taken against using generic MSK models for personalized JRFs estimation, especially when the bony torsional profile differs significantly from the generic model. Further studies are required to confirm these hypotheses and to determine whether altering the model musculoskeletal anatomy requires adjustments in muscle architectural parameters. The presented methodology and bone-deformation tool can be easily applied to investigate the musculotendon kinematics in the presence of distal femoral or tibial torsions (see Figure S1 of supplementary materials for examples) or extended to study the dependency of JRFs on bone morphology in other human joints.

## Appendix

To facilitate the reproducibility and replication of our results, we have released our research code and data with this publication. All of the data and scripts needed to run the calculations reported in this work, as well as the post-processing scripts to reproduce the figures in the paper are available at https://github.com/modenaxe/femoral_anteversion_paper.

## CRediT authorship contribution statement

**Luca Modenese:** Conceptualization, Methodology, Software, Data Curation, Validation, Visualization, Formal analysis, Writing – original draft.

**Martina Barzan:** Conceptualization, Software, Writing - review & editing.

**Christopher Carty:** Conceptualization, Writing - review & editing.

## Acknowledgements

LM was supported by an Imperial College Research Fellowship granted by Imperial College London and wants to thank Arnault Caillet and Davide Conte for the feedback on the manuscript. CC and MB were supported by a Queensland Advancing Clinical Research Fellowship granted by the Queensland Government. MB was supported by an ARC ITTC grant (ARC CMIT) from the Australian Government.

