## supplementary materials for "Dependency of Lower Limb Joint Reaction Forces on Femoral Anteversion"

### Web Supplementary materials

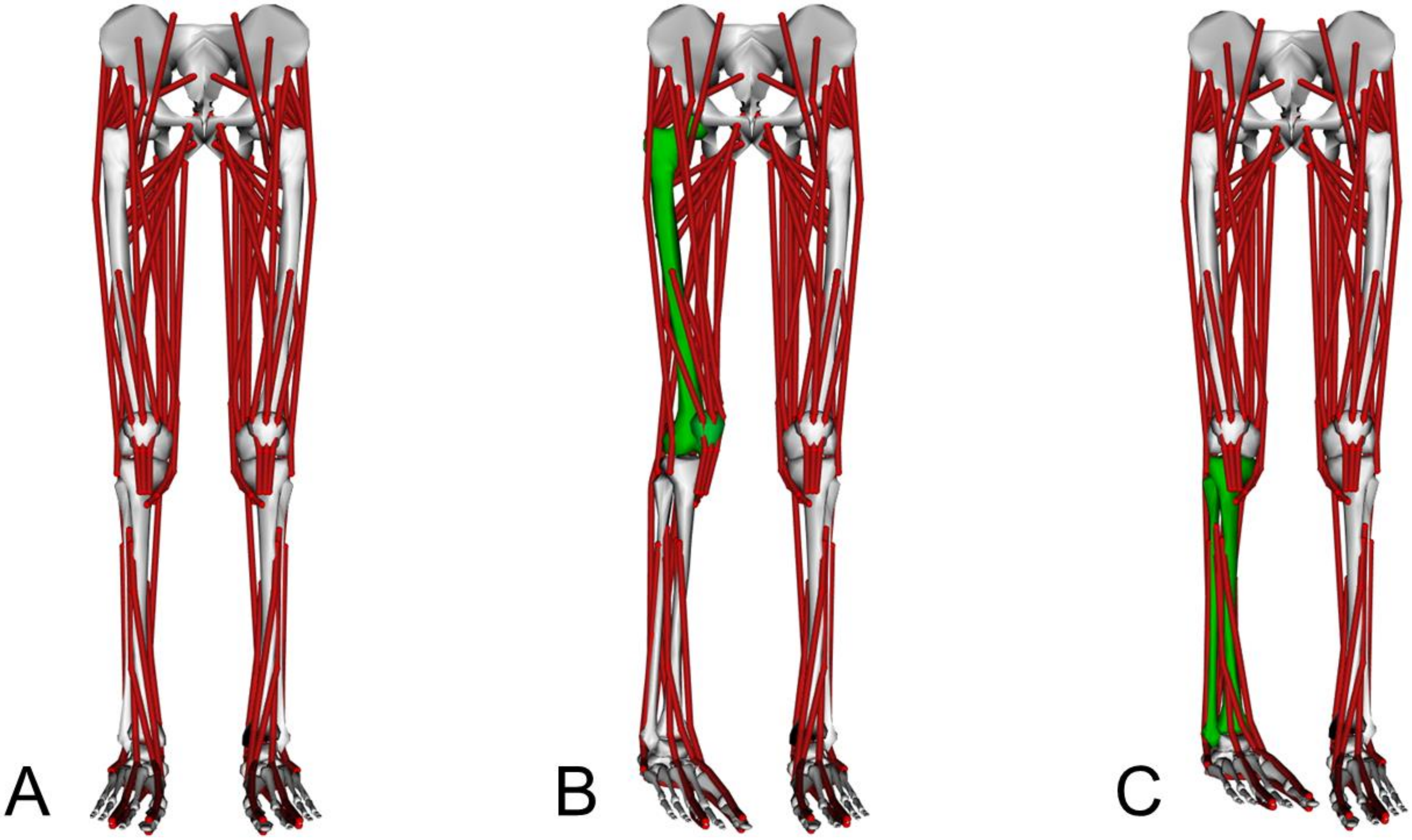


Figure S1 Possible model alterations enabled by the bone-deformation tool described in the main manuscript when the joint parameters are modified together with the bone morphology: the generic musculoskeletal model of Rajagopal et al. (2016) (A) can be modified to represent a distal femoral torsion (B), or a distal tibial torsion (C) affecting the foot alignment. Modified bones are colored green and both applied torsions are 30˚. Note how model segments connected to the deformed bone, e.g. the patella in the case (B), are also modified automatically to maintain consistency with the deformed model.

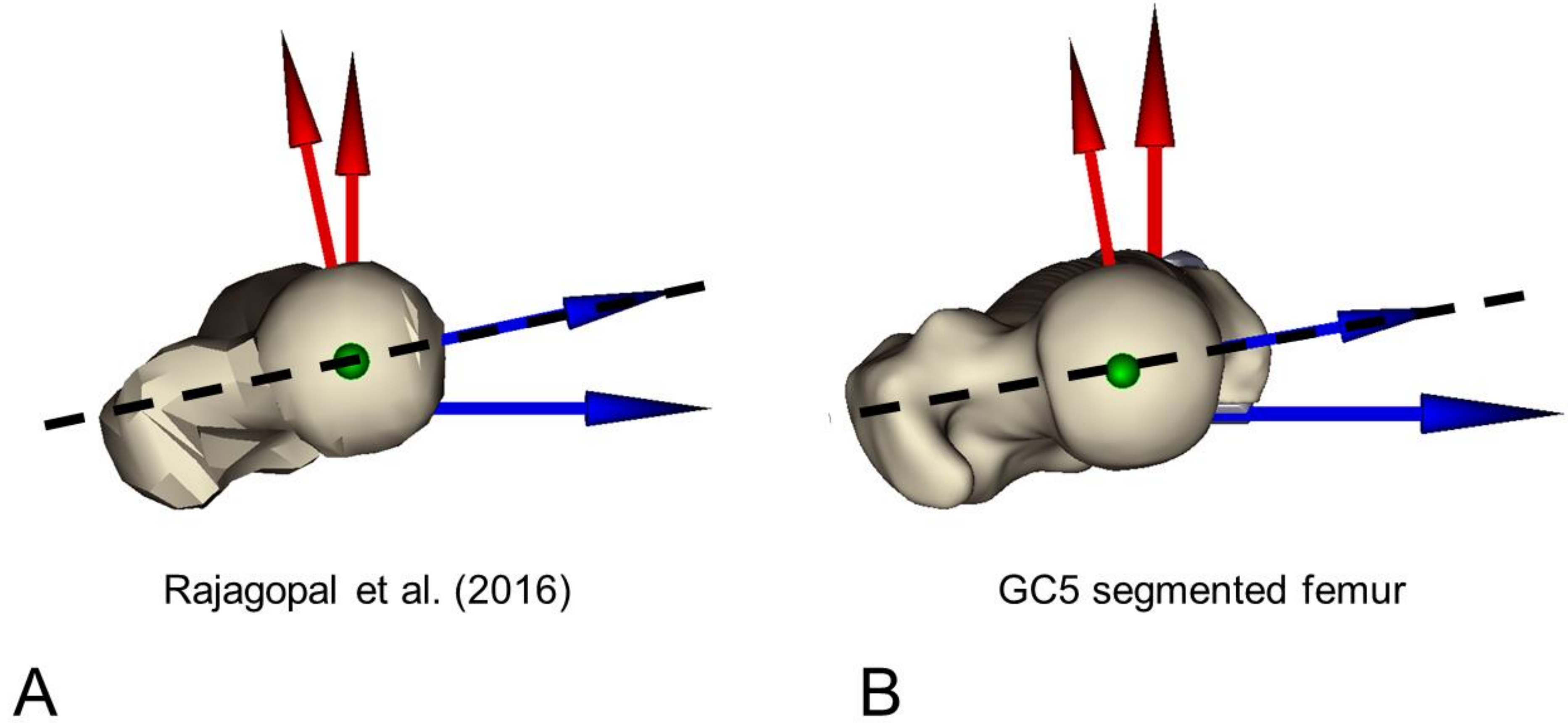


Figure S2 Estimation of the femoral anteversion angles in the femoral geometries of the baseline model (around 12˚) (A) and on the segmented femoral geometry available on the 5th Grand Challenge dataset (around 10˚) (B). Note the low quality of femoral bone surface provided with the musculoskeletal model. Angles were estimated using NMSBuilder (Valente et al., 2017), aligning the Z axis of the proximal reference system (in blue) with the femoral neck from a proximal view as in the figure and quantifying the angle with respect to the Z axis of the distal reference system (aligned with the posterior condylar axis).
